# Hydration shells of globular and intrinsically disordered proteins: effects of amino acid composition, peptide conformation, and force fields

**DOI:** 10.1101/2025.06.10.658889

**Authors:** Johanna-Barbara Linse, Tobias M. Fischbach, Jochen S. Hub

## Abstract

The protein hydration shell is a key mediator of processes such as molecular recognition, protein folding, and proton transfer. How surface-exposed amino acids shape the hydration shell structure is not well understood. We combine molecular dynamics simulations with explicit-solvent predictions of small-angle X-ray scattering (SAXS) curves to quantify the contributions of all 20 proteinogenic amino acids to the hydration shell of the globular GB3 domain and the intrinsically disordered protein (IDP) XAO. We focus on two quantities encoded by SAXS curves: the hydration shell effect on the radius of gyration and the electron density contrast between protein and solvent. We derive an amino-acid-specific contrast score, revealing that acidic residues generate the strongest contrast with 1 to 1.5 excess water molecules relative to alanine, followed by cationic and polar residues. In contrast, apolar residues generate a water depletion layer. These trends are consistent across simulations with different water models. Around the XAO peptide, the hydration shell is generally far weaker compared to the globular GB3 domain, indicating unfavorable water–peptide packing at the IDP surface. The hydration shell effect on the radius of gyration of the IDP is strongly conformation-dependent. Together, the calculations show that the composition and spatial arrangement of surface-exposed amino acids govern the hydration shell structure, with implications for a wide range of biological functions and for hydration-sensitive experimental techniques such as solution scattering.

**Significance:** Hydration shells of biomolecules constitute a large fraction of the water in crowded cellular environments and play key roles in biological functions such as enzymatic reactions and conformational transitions. Small-angle X-ray scattering (SAXS) has shown that hydration shells differ in density from bulk water, yet how surface-exposed amino acids and protein surface geometry shape the hydration shell is not well understood. We combined molecular dynamics simulations with explicit-solvent SAXS predictions to quantify how surface-exposed chemical moieties and protein geometry drive variations in hydration shell density. Notably, the hydration shell of a globular protein differs markedly from that of an intrinsically disordered protein. Our study offers a comprehensive characterization of protein hydration and informs the interpretation of hydration-sensitive experimental techniques.

## Introduction

Proteins in solution are enveloped by a hydration shell, formed through electrostatic and dispersive interactions between water molecules and surface-exposed protein moieties. The hydration shell actively participates in various biological functions such as protein folding, molecular recognition, enzyme catalysis, proton transfer, or avoidance of unspecific aggregation, and is thus considered an integral part of proteins.^1–5^ The structure and dynamics of the hydration shell differ from those of bulk water, as revealed by nuclear magnetic resonance, terahertz spectroscopy, time-dependent fluorescence Stokes shift, inelastic neutron scattering, molecular dynamics (MD) simulations, and several other techniques.^6–16^ Consequently, the vibrational, rotational, and translational dynamics of water molecules in the hydration shell are slowed down by approximately two-to fivefold. Scattering experiments have revealed that the water density in the hydration shell of many proteins is increased relative to the density of bulk water, with the magnitude of this density increase likely being protein-dependent.^17–20^ In the crowded cytoplasm of biological cells, up to 70% of water belongs to a hydration shell, indicating that water involved in life is predominantly non-bulk-like.^3,21,22^ How the composition and relative arrangement of surface-exposed amino acids control the properties of the protein hydration shell is not well understood. Small-angle scattering with X-rays and neutrons (SAXS/SANS) of highly charged proteins suggested that surface-exposed anionic aspartate or glutamate residues increase the hydration shell density more than cationic lysine or arginine residues,^20,23^ which aligns with the large number of structured water located at anionic residues in protein crystals.^24^ Furthermore, spectroscopic techniques revealed that the polarity of surfaces influences the properties of water at interfaces. At polar surfaces, water exhibits decreased internal water order and fewer internal hydrogen bonds. At apolar surfaces, in contrast, water exhibits increased internal order and more internal hydrogen bonds, and it may form low-density clathrate structures.^25–35^ Additionally, water has been shown to form a depletion layer with reduced density at hydrophobic surfaces. ^36–38^ However, the quantitative influence of surface-exposed amino acids or of specific chemical moieties on the hydration shell architecture remains largely unexplored.

We recently validated the protein hydration shell from MD simulations by comparing results from explicit-solvent SAXS/SANS predictions^39–41^ with consensus experimental data obtained from a worldwide community effort.^20,42^ SAXS and SANS data reflect the contrast of the, respectively, electron density or neutron scattering length density of the protein relative to bulk solvent, thereby including contributions of the hydration shell. We observed that many but not all combinations of protein force fields and water models accurately reproduce the hydration shell effect on the radius of gyration *R*_g_. We furthermore found that the hydration shell effect on *R*_g_ depends on protein size, geometry, and surface composition, suggesting that the effect represents a protein-specific footprint of the hydration shell. In this study, we use MD simulations to quantify the influence of all proteinogenic amino acids on the hydration shell of a globular and an intrinsically disordered protein on two parameters that are encoded by SAXS curves, namely on the *R*_g_ and on the overall contrast between solute and solvent. We derive an amino-acid-specific contrast score for solvent-exposed proteinogenic residues and show that the hydration shell structure and its effect on SAXS data depends not only on chemical composition but also on peptide conformation and water models.

## Results

To quantify the effects of amino acid composition on the hydration shell of proteins, we simulated the GB3 domain as a representative for globular proteins (Fig. 1A/B) and the XAO peptide^43,44^ as representative for IDPs (Fig. 1C/D). The three-dimensional electron density of solvent around the GB3 domain or around XAO are shown in Figs. 1B/D, here computed from simulations with position restraints on all heavy atoms leading to spatially well-defined densities from surface-bound water molecules. The densities reveal highly localized water molecules (red densities), the first hydration shell (orange/red densities) as well as the second hydration shell (dark blue densities). A highly shallow third shell is hardly visible in the three-dimensional density representation (Figs. 1D, cyan density layer). The structure of the hydration shell involving a pronounced first shell, a shallow second, and a highly shallow third shell agrees with many previous MD studies (Ref. 45 and references therein).

**Figure 1.**
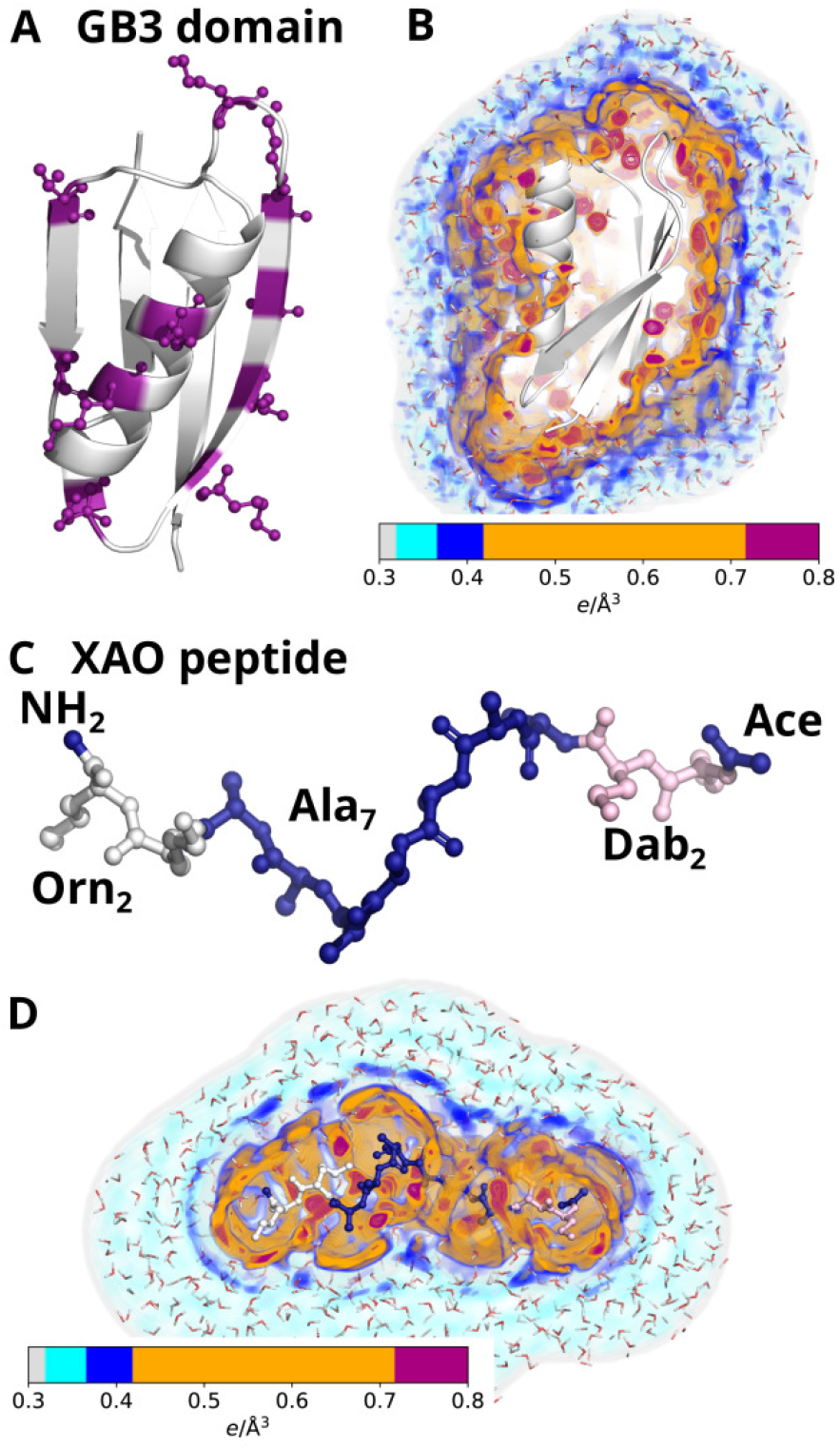
(A) Cartoon representation of the GB3 domain. Ten surface amino acids shown in purple ball-and-stick representation were mutated into each of the 20 proteinogenic amino acids. (B) Three-dimensional density of the hydration shell around the wild type GB3 domain (for colors see colorbar). The first and second hydration layers appear as orange and blue densities, respectively. Red densities indicate well-defined water position at the protein surface. (C) XAO peptide with four unnatural amino acids at the termini: two 2,4-diaminobutyric acid (Dab) and two ornithin (Orn) shown in pink and white, respectively. (D) Three-dimensional solvent density around the wild type XAO peptide.

### The hydration shell of the globular GB3 domain strongly depends on the surface amino acid composition

SAXS experiments of proteins probe the electron density contrast between the protein and the bulk solvent, including the density contrast contributed by the hydration shell. To quantify the effects of different amino acids to the hydration shell, and to relate variations among different amino acids to putative solution scattering experiments, we computed SAXS curves of the wild type and of 21 mutants of the globular GB3 domain. We selected ten surface-exposed residues (Fig. 1A, pink ball-and-stick representation) and mutated these residues to each of the 20 proteinogenic amino acids, while including histidine in the neutral form (*δ*-nitrogen protonated, His^0^) and in the cationic form (*δ*-and *ϵ*-nitrogen protonated, His^+^), resulting in 22 GB3 variants (wild type and 21 mutants). We performed explicit-solvent MD simulations with restraints on the backbone atoms to maintain all GB3 variants in identical backbone conformation and to prevent unfolding of putatively unstable GB3 mutants such as mutants with many hydrophobic surface-exposed residues. SAXS curves *I*(*q*) computed for the 22 GB3 variants differ (Fig. 2). Since we computed the SAXS curves taking the solvent explicitly into account,^39,41,46^ the variations among the SAXS curves include effects owing to variations of the hydration shell contrast.

**Figure 2.**
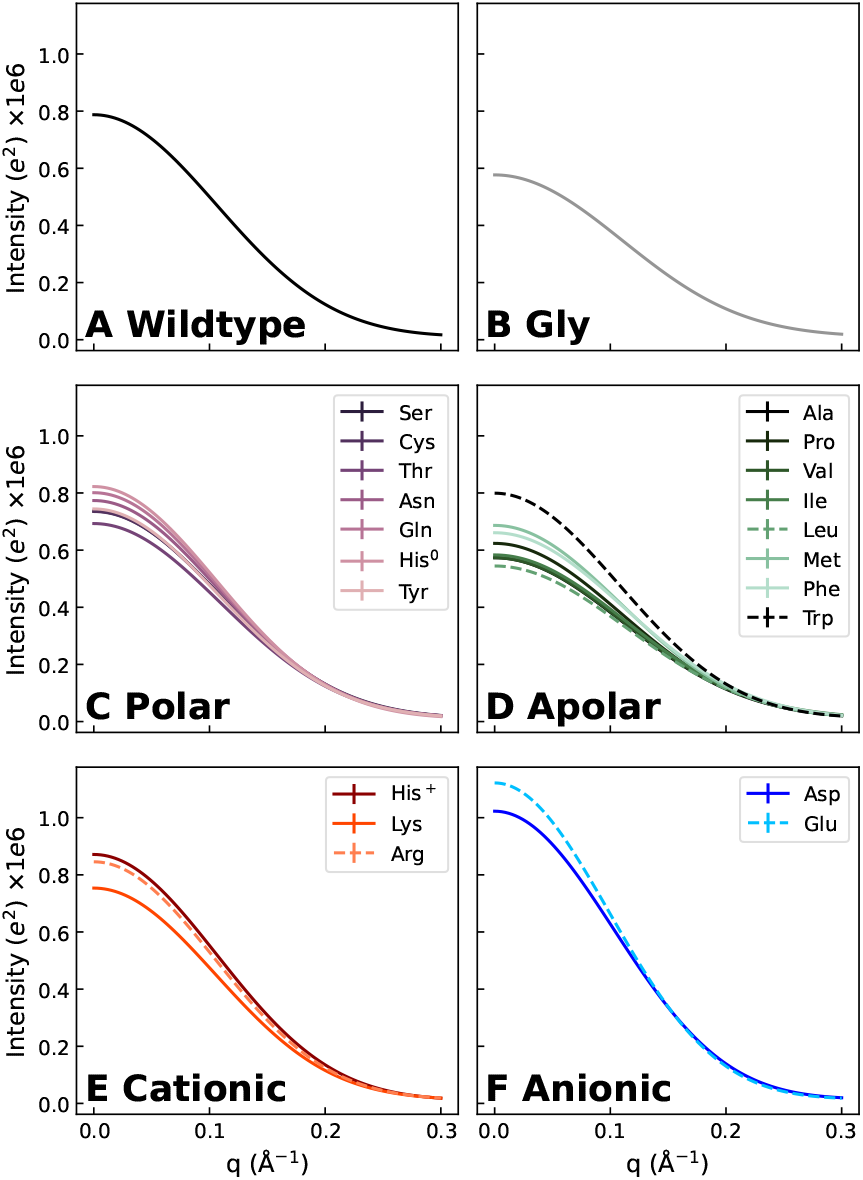
SAXS curves of the GB3 domain from explicit-solvent SAXS calculations with the TIP4P/2005 water model in combination with the ff03w protein force field (A) for GB3 wild type or (B–F) for 21 GB3 variants with 10 mutated surface-exposed amino acids each (for color code and line style, see legends). For clarity, SAXS curves are grouped by the amino acid property (glycine, polar, apolar, cationic, anionic) in panels B–F.

In this study, we used two quantities that provide a footprint for the hydration shell while being encoded by the SAXS curves: (i) the forward scattering intensity *I*_0_ = *I*(*q* = 0), which is related to the contrast between protein and solvent; and (ii) the radius of gyration *R*_*g*_, which quantifies the spatial extent of the protein. Focusing first on the former quantity, the forward scattering is given by the square of the total contrast Δ*N*_*e*_ in number of electrons between protein and solvent, i.e.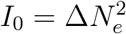. We decomposed the total contrast into contributions from the contrast of the bare protein 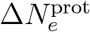 and the contrast of the hydration shell 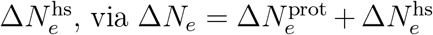 (see Methods). Figure 3A (yellow bars) presents the contrast of the hydration shell for 22 GB3 variants, here computed with the Amber force field ff03w in conjunction with the TIP4P/2005 water model,^47,48^ which revealed excellent agreement with experimental SAXS/SANS data in our previous study and may, therefore, be taken as reference force field. ^20^ The contrast is plotted in number of water molecules as 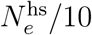 given that each water molecule contains 10 electrons. Evidently, the contrast of the hydration shell differs greatly among GB3 variants with different surface-exposed amino acid types. The hydration shell of wild type GB3 exhibits a positive contrast of 4.1 water molecules implying the presence of a additional 4.1 water molecules in the hydration shell relative to an equivalent volume of bulk water, in line with the well-known densely packed hydration shell documented by SAXS experiments of several proteins.^17,18,20^ Among the mutated GB3 variants, the variants with additional anionic residues (Asp/Glu) reveal the largest contrast, followed in decreasing order by GB3 variants with cationic (Lys/Arg/His^+^), and polar charge-neutral amino acids (Fig. 3, pink labels at the abscissa). The marked hydration shell imposed by the anionic residues Glu/Asp aligns with previous SAXS/SANS experiments of super-charged variants of green fluorescent protein^49^ and of the highly anionic glucose isomerase.^20,42^ In contrast, GB3 variants with many apolar surface-exposed residues reveal a negative hydration shell contrast, indicating a water depletion layer in the vicinity of hydrophobic amino acids, in line with reports for other types of hydrophobic surfaces.^36–38^ An exception to the order anionic–cationic–polar–apolar is given by tyrosine, which displays a more negative contrast compared to all other polar amino acids, rationalized by the presence of the apolar six-membered benzene ring.

**Figure 3.**
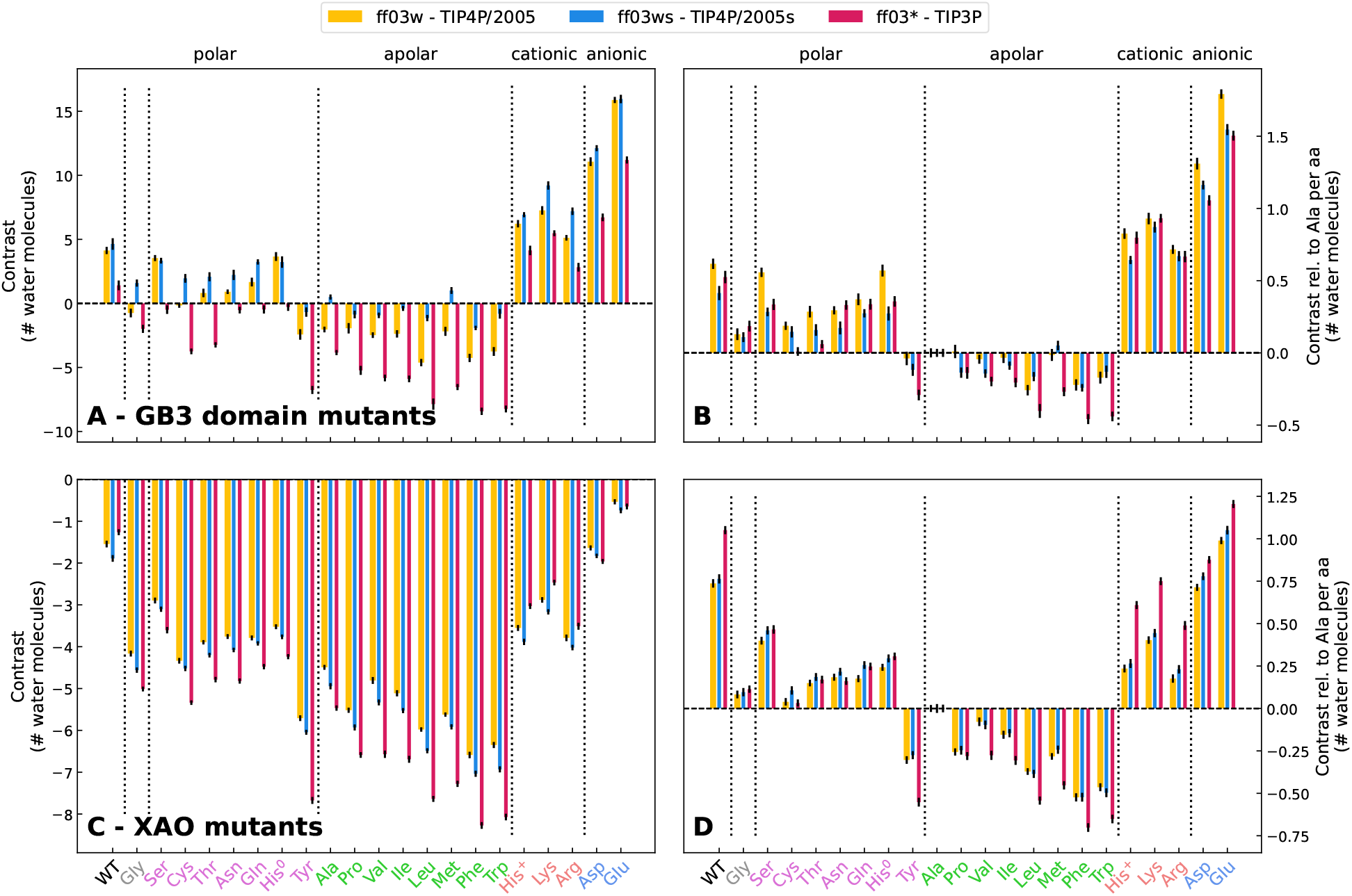
(A) Contrast of the hydration shell in number of water molecules of GB3 wild type and 21 GB3 mutants, see labels at abscissa colored by the property of the amino acid: Gly (grey), polar (pink), apolar (green), cationic (orange), anionic (blue) residues. Contrast values are shown for three combinations of protein force field and water model: ff03w–TIP4P/2005 (yellow), ff03ws–TIP4P/2005s (blue), ff03^∗^–TIP3P (red). (B) Contrast per amino acid for GB3 domain relative to alanine. (C/D) Same analysis as in panels (A/B) for the XAO peptide.

To quantify the hydration shell contrast imposed by individual amino acids, Fig. 3B presents the contrast per amino acid and relative to alanine as reference. Accordingly, the hydration shells of anionic residues exhibit approximately 1.5 additional water molecules relative to alanine, with glutamate standing out as the amino acid whose hydration shell imposes the largest contrast, indicative for a particularly densely packed hydration shell. The hydration shells of cationic residues contain roughly one additional water molecule, whereas the hydration shells of polar residues contain approximately 0.2 to 0.5 additional water molecules relative to alanine. Bulky apolar amino acids such as leucine or phenylalanine may contain up to 0.3 fewer water molecules relative to alanine. Tyrosine with its polar hydroxyl group and apolar aromatic ring represents an intermediate case between polar and apolar residues. These values provide an amino acid-resolved hydration layer contrast score for a common globular protein such as GB3, thus quantifying how chemical specificities of proteinogenic amino acids control the density of the protein hydration shell. Below we further analyze these data to dissect how individual chemical moieties control the hydration shell.

### Among different water models, effects of amino acid properties on the hydration layer contrast agree qualitatively but differ quantitatively

To test the influence of water models, and to exclude that our key findings are not biased by the choice of the water model, we calculated the contrasts for three different water models, namely TIP3P,^50^ TIP4P/2005,^48^ and TIP4P/2005s,^51^ in combination with their corresponding Amber03 force field^52^ variant. Among these water models, TIP3P is the most widely used model since the popular Charmm and Amber protein force fields have originally been parameterized in conjunction with TIP3P. However, TIP3P shows poor agreement with experimental data as it yields a too low density, a too high diffusion coefficient, and a too high isothermal compressibility.^53^ TIP4P/2005 reproduces water properties more accurately as compared to TIP3P, and it furthermore captures the water density over a wide temperature range.^48^ TIP4P/2005s takes water–water interactions from TIP4P/2005, however it implements increased water–protein dispersion interactions with the aim to balance water–water against water–protein interactions in protein simulations with Amber force fields.^51^

Figure 3A compares the total hydration shell contrast of 22 GB3 variants for simulations with these three water models (yellow, blue, red bars, see legend). Irrespective of the water model, the contrasts follow the order anionic–cationic–polar–apolar, suggesting that the effects of amino acid classes on the hydration shell contrast agree qualitatively among the water models. This finding is confirmed by comparing the contrast per amino acid relative to alanine (Fig. 3B). However, the total contrasts shown in Fig. 3A furthermore reveal considerable quantitative differences among water models. TIP3P yields the lowest contrast for all GB3 variants with approximately four water molecules fewer within the overall GB3 hydration shell compared to TIP4P/2005 (Fig. 3A, red vs. yellow bars). In contrast, TIP4P/2005s yields an increased contrast for most GB3 variants with typically zero to three additional water molecules in the hydration shell compared to TIP4P/2005 (blue vs. yellow bars), rationalized by the increased protein–water dispersion interactions implemented by TIP4P/2005s. ^51^ More specifically, TIP4P/2005s yields a similar contrast compared to TIP4P/2005 for the GB3 wild type and for the Ser, His^0^, His^+^, Asp, Glu, and an increased contrast for all the other variants. Thereby, the hydration shell contrast by TIP4P/2005s largely exceeds the contrast by TIP3P, in particular for the apolar variants, with up to 7.5 additional water molecules in the hydration shell for the Trp variant.

### Hydration shell contrasts controlled by amino acid type and force field impose experimentally accessible variations of the radius of gyration

Analysis of the hydration shell based on *I*_0_ and hydration shell contrasts involve two caveats. First, experimental *I*_0_ values –or, equivalently, total electron density contrasts– are subject to relatively high uncertainty because obtaining *I*_0_ from an experimental SAXS curve would require precise knowledge of the solute concentration. Because the solute concentration is typically only approximately known, quantitative comparisons of *I*_0_ between MD simulations and experiments is difficult. In contrast, the radius of gyration *R*_g_ obtained by SAXS coupled to size exclusion chromatography (SEC-SAXS) enables *R*_g_ measurements with sub-Ångström accuracy,^42^ thereby enabling quantitative validation of MD simulations against experiments.^20^ Second, whereas the overall solute contrast is unambiguously defined in explicit-solvent SAXS calculations via (*I*_0_)^1*/*2^, its decomposition into contrast contributions from the protein and hydration shell depends on the definition of the protein volume or, equivalently, on the definition of the dividing surface at the protein–water interface.^45^ In contrast, computing the hydration shell effect on *R*_g_ does not require a definition of the dividing surface.

Thus, as a second indicator for the hydration shell, we analyzed hydration shell effects on the *R*_*g*_ values of 22 GB3 variants. We computed the change of *R*_*g*_ owing to the hydration shell, as given by the 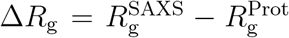, where 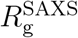 denotes the *R*_g_ value obtained by Guinier analysis of the SAXS curve, thereby taking the hydration shell into account, and 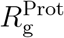 denotes the *R*_g_ value of the bare proteins computed from the atomic coordinates of protein atoms. Figure 4A presents Δ*R*_g_ values for the 22 GB3 variants, whereas Fig. 4B shows the Δ*R*_g_ values relative to alanine per mutated amino acid. In line with the contrasts discussed above, the presence of anionic residues (Asp/Glu) impose by far the largest increase of Δ*R*_g_ values by ∼1.5 Å, and an increase per amino acid relative to alanine by ∼0.1 Å. These results confirm that anionic residues impose a tightly packed hydration shell.^20,24,49^ In contrast, bulky hydrophobic residues such as Val, Ile, Leu, Phe, or Trp lead to small Δ*R*_g_ values, and a decrease per amino acid relative to alanine by up to ∼0.05 Å, in line with a water depletion layer at a the hydrophobic surface. Cationic and many polar residues lead to intermediate and similar Δ*R*_g_ values, which may be surprising considering that hydration shell contrasts imposed by cationic residues clearly exceed the contrasts imposed by polar residues (see Fig. 3A/B). They findings might reflect that Δ*R*_g_ is sensitive to the spatial distributions of contrasts, which may lead to different amino acid-specific effects on Δ*R*_g_ compared to effects on the total contrast derived from *I*_0_.

**Figure 4.**
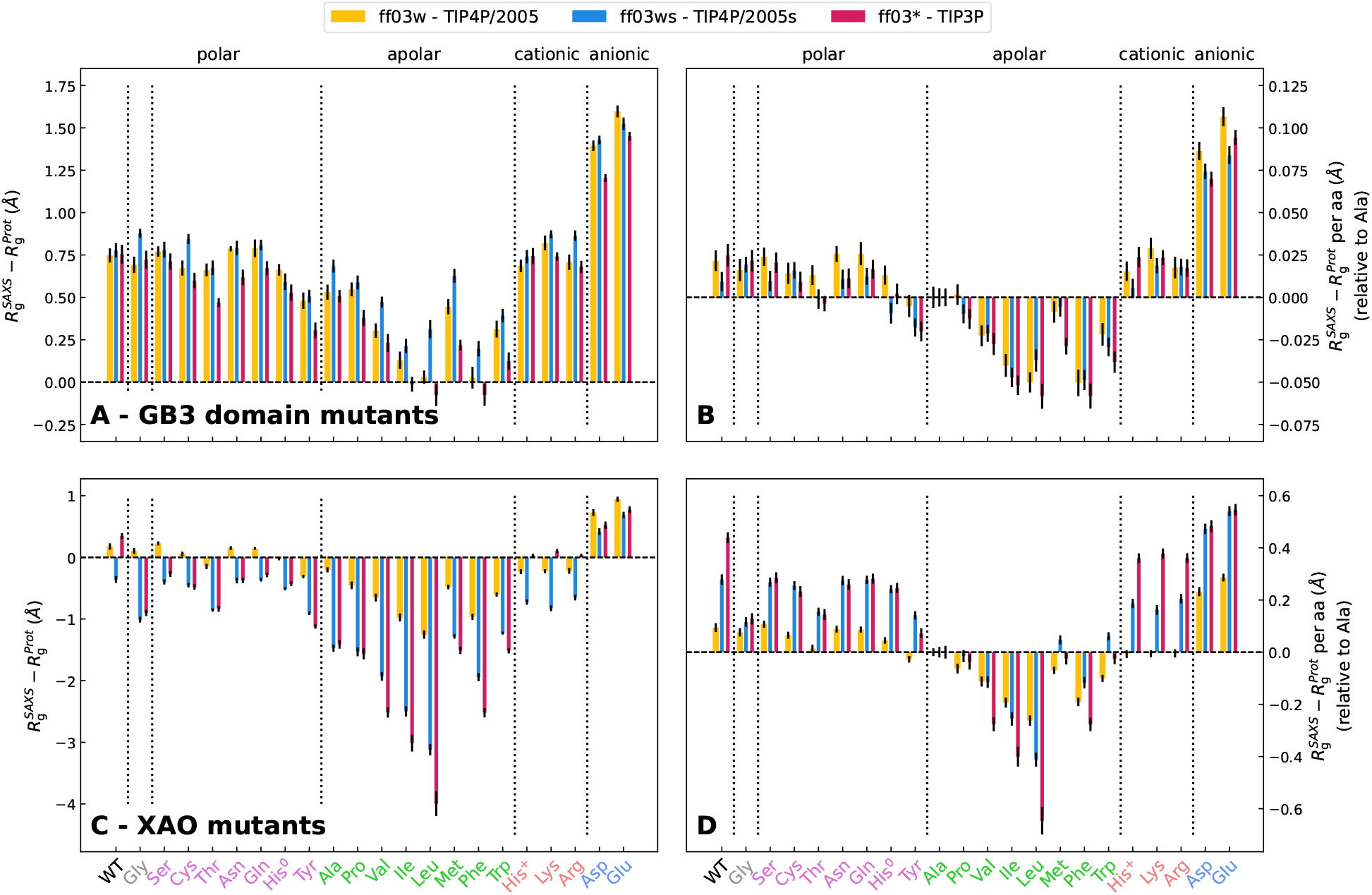
(A) Hydration-induced shift in the radius of gyration Δ*R*_g_ from explicit-solvent SAXS calculation of 22 GB3 variants. Δ*R*_g_ values are shown for GB3 wild type (WT) and 21 mutants, see labels at the abscissa colored by the property of the amino acid: Gly (grey), polar (pink), apolar (green), cationic (orange), anionic (blue) residues. Results are shown for three combinations of protein force field and water model: ff03w–TIP4P/2005 (yellow), ff03ws–TIP4P/2005s (blue), ff03^∗^–TIP3P (red). (B) Δ*R*_g_ values relative to the alanine mutant and per amino acid. (C/D) Same analysis as in panels (A/B) for the XAO peptide.

Comparing the Δ*R*_g_ values between simulations with TIP3, TIP4P/2005, and TIP4P/2005s reveals specific properties of water models, although the relative differences among Δ*R*_g_ values are overall smaller as compared to the differences among the contrasts (compare Fig. 4A/B with Fig. 3A/B). Whereas the Δ*R*_g_ values of GB3 wild type is in excellent agreement among the three water models (4A, left column), TIP3 yields lower Δ*R*_g_ for many GB3 mutants such as mutants with anionic residues (Asp/Glu), several apolar residues, as well as for several apolar residues such as Thr or Tyr. The largest variations of Δ*R*_g_ values among water models is found for the bulky apolar residues, where in particular TIP4P/2005s, but also TIP4P/2005, yields by far larger Δ*R*_g_ values as compared to TIP3P. Thus, the increased protein–water dispersion interactions implemented by TIP4P/2005s leads to a partial loss of the water depletion layer at hydrophobic surfaces, with a footprint on the radii of gyration.

### Hydration shell of the intrinsically disordered protein (IDP) XAO exhibits a negative amino acid-dependent contrast

Compared to globular proteins, IDPs exhibit a larger surface-to-volume ratio, suggesting that –at a given protein density– IDPs exhibit more protein–water contacts and perturb a larger volume of water, rationalizing the tight coupling between IDP and water dynamics.^54^ Many IDPs carry out their function by partial folding on the surface of other proteins, thereby involving large rearrangements of protein–water and water–water interaction networks. Nevertheless, the hydration shell of IDPs has attracted less attention compared to the hydration shell of globular proteins.^55,56^ A previous MD study suggested that accurate representation of the hydration shell density by explicit solvent models is critical for predicting SAXS curves of IDPs since even small variations of the hydration shell density may strongly influence predicted SAXS curves.^19,57^ MD simulations using TIP4P/2005 or TIP4P/2005s revealed that the water structure in the hydration shell of an IDP is perturbed relative to the bulk as indicated by a loss of tetrahedrality, however this perturbation has been weaker as compared to the hydration shell of a folded protein.^58^ How the amino acid composition of an IDP controls the density of its hydration shell has not been systematically addressed.

As a model IDP, we here consider the XAO peptide with the sequence Ace-(diaminobutyric acid)_2_-(Ala)_7_-(ornithine)_2_-NH_2_, whose conformational ensemble has been studied by SAXS as well as by NMR and circular dichroism spectroscopy.^43,44^ We used maximum-entropy ensemble refinement against SAXS data taken from Ref. 43 to obtain the heterogeneous ensemble of XAO. We randomly selected 20 frames from the ensemble, thereby representing the heterogeneous ensemble, involving compact and expanded XAO conformations (see Methods and Fig. S1). In follow-up simulations of XAO and its mutants, these conformations were maintained using backbone restraints.

To reveal how the amino acid composition influences the hydration shell of an IDP, we mutated the four terminal XAO residues to 20 proteinogenic amino acids, again including histidine in the neutral and cationic form (21 mutants). For each XAO variant, we carried out 20 MD simulations with backbone restraints to the 20 conformations taken from the XAO ensemble (see above) and computed the SAXS curve using explicit solvent SAXS calculations. Following the analysis described above for the GB3 domain, we obtained the contrast of the hydration the XAO wild type and 21 mutants (22 variants), using the three water models TIP4P/2005, TIP4P/2005s, and TIP3P. We computed the overall contrast of the hydration shell relative to an equivalent volume of bulk water (Fig. 3C) as well as the contrast per amino acid relative to alanine (Fig. 3D).

At variance with the analysis for GB3, we found a negative hydration shell contrast for all 22 XAO variants, implying that the hydration shell of XAO contains fewer water molecules as compared to bulk water. Whereas XAO wild type and anionic mutants reveal a contrast of up to approx. −2 water molecules, apolar variants may reveal contrast of up to −7 water molecules or even less (Fig. 3C). We rationalize the negative contrasts with the high flexibility of XAO, leading to more loosely packed conformations as compared to the structure of the globular GB3. Thereby, small voids between the XAO backbone and side chains may exclude water molecules or enable only incomplete packing of water around XAO moieties, leading to a lower water density around XAO as compared to GB3.

The effects of amino acid classes on the hydration shells are consistent with the findings for GB3, namely the contrasts follow roughly the order anionic–cationic–polar–apolar (Fig. 3D). However, contrasts per amino acid differ quantitatively for XAO compared to our results for GB3. For instance, arginine residues imposed smaller contrasts in XAO compared to arginine in GB3 (compared Fig. 3B with D), possibly because the hydrophobic C_*β*_, C_*γ*_, and C_*δ*_ atoms of arginine near the XAO termini are more solvent-exposed compared to arginines at the GB3 surface.

### The hydration shell of an IDP may reduce the radius of gyration detected by SAXS

In sharp contrast to the findings for GB3 variants, for which Δ*R*_g_ values were mostly positive in the range of approximately 0 to 1.5 Å, Δ*R*_g_ values for XAO variants with different force fields are mostly negative and may take large absolute values (Fig. 4C). The radius of gyration detected by SAXS is given by

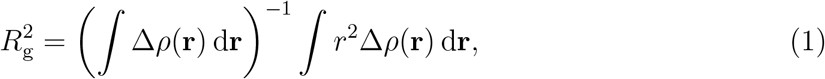

where Δ*ρ*(**r**) denotes the electron density contrast and *r* the distance from the (contrast-weighted) center of mass. The large negative Δ*R*_g_ values up to −4 Å may be rationalized by (i) the overall small contrast of XAO (small value in brackets in Eq. 1), leading to a large impact of the hydration shell contrast on *R*_g_, and (ii) hydration shell contributions to the contrast close to the center of mass (at small *r*), as occurring for extended conformations for which moieties near the center of mass are solvent-exposed. Positive Δ*R*_g_ values are found only for the anionic mutants for which Asp/Glu residues may impose large contrast near the endpoints of XAO, but also for the XAO wild type and few polar variants, mostly with the TIP4P/2005 water model (Fig. 4C, yellow bars). Thus, whereas the hydration shell of globular proteins often leads to an increased *R*_g_ detected by SAXS,^17,23,39^ the hydration shell of IDPs may also lower the *R*_g_ value.

### Three-dimensional densities around XAO reveal amino-acid and force field-specific hydration shell structures

To rationalize the variations of contrast and *R*_g_ values in structural terms, we computed the three-dimensional solvent densities with the TIP4P/2005 water model around extended conformations of four XAO variants, in which the four terminal residues were mutated to serine, leucine, lysine, or glutamate, providing one representative each for a polar, apolar, cationic, or anionic variant (Fig. 5E–H). In line with the findings from the XAO ensemble (Fig. 3C/D and 4C/D), these mutations lead to large variations of the forward scattering *I*_0_ (Fig. 5A), contrast of the hydration shell between approximately −1 and −6 water molecules (Fig. 5B), *R*_g_ values obtained from SAXS between ∼7.5 Å and ∼11.5 Å (Fig. 5C, blue squares), and Δ*R*_g_ values between −3 Å and +1 Å (Fig. 5D). These value align qualitatively with the variations of solvent densities near the endpoints of XAO. For instance, the solvent structure around the anionic glutamate residues reveals several tightly bound water molecules (Fig. 5H, red spots) and a pronounced second hydration shell. In contrast, the hydration shell around the apolar leucine lacks any structured water molecules, while the second hydration shell is blurred out (Fig. 5F). The serine and lysine variants yield intermediate cases (Fig. 5E/G). Thus the variations of contrast and *R*_g_ values quantified above are footprints of amino acid-specific hydration shell structures revealed by three-dimensional solvent densities.

**Figure 5.**
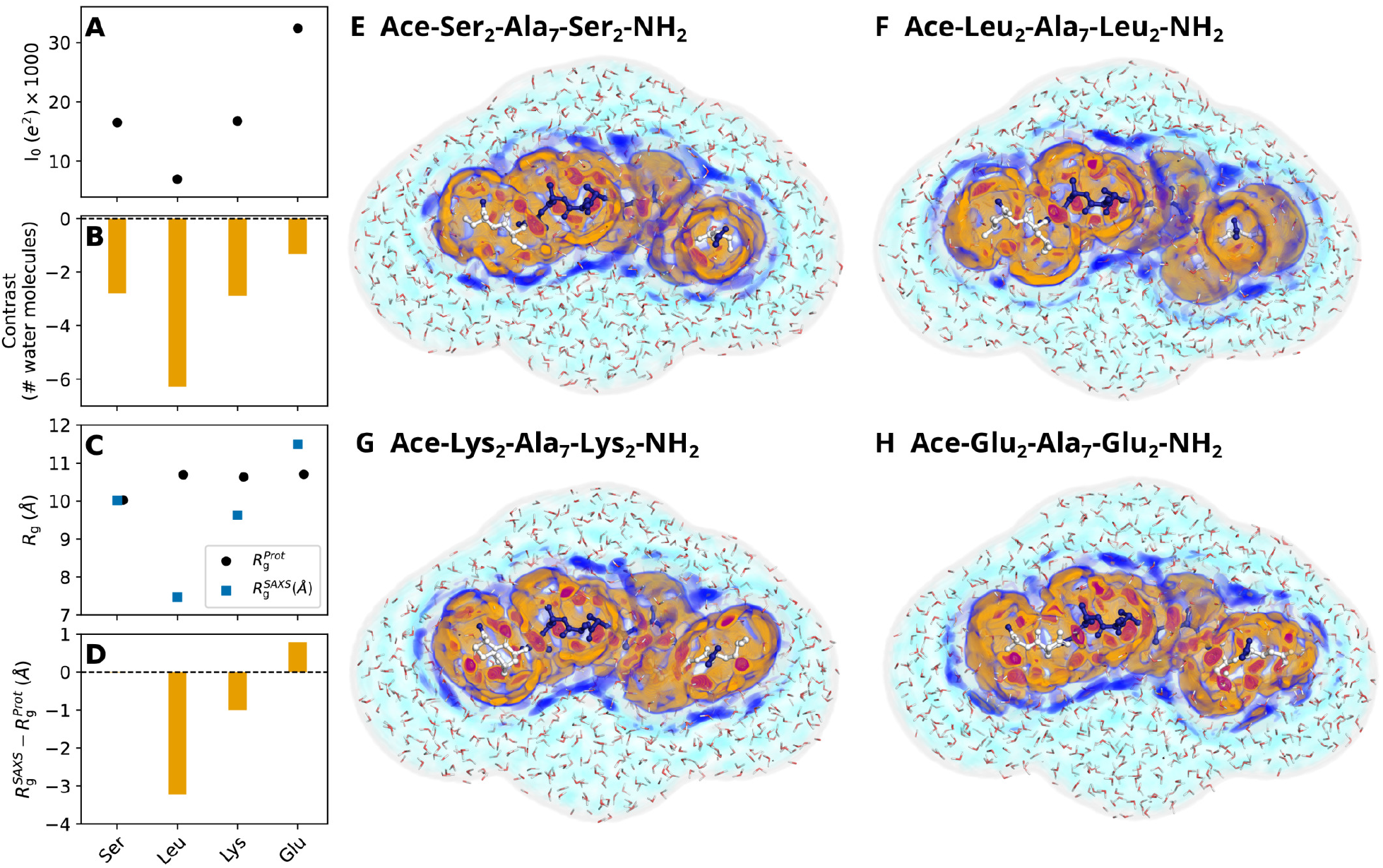
Comparison of hydration shells of XAO mutants with different types of mutated residues: Ser (polar), Leu (apolar), Lys (cationic), Glu (anionic). (A) Forward scattering *I*_0_, (B) hydration shell contrast in number of water molecules, (C) radius of gyration 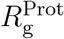 and 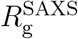 from the bare peptide (black dots) and *R*^SAXS^ from Guinier analysis (blue squares), and (D) 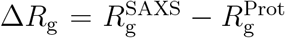. Values were calculated from simulations with the TIP4P/2005 water model in combination with the ff03w protein force field. (E–H) Shaded colors show the three-dimensional solvent density maps from 50 ns simulations around the mutants analyzed in panels A–D with color code taken from Fig. 1B/D. MD simulation used to compute density maps were carried out with restraints on all heavy atoms, thereby yielding spatially well-defined hydration shells. The density is overlayed with one MD frame, showing the XAO in ball-and-stick representation, and water molecules within the envelope as red/white lines. Color code according to Fig.1B/D.

The solvent densities around the extended XAO peptide depends not only on amino acid composition but furthermore on the water model, as evident, for instance, from more pronounced solvent structures modeled by TIP4P/2005s as compared to TIP3P (Fig. S3). Such force field-dependent solvent densities rationalize variations of hydration shell contrasts by the ensembles of 22 XAO variants presented in Fig. 3C/D, which qualitatively align with the findings for GB3 (Fig. 3A/B). Specifically, TIP3P yields smaller (more negative) hydration shell contrasts relative to TIP4P/2005, whereas the increased protein–water dispersion interactions implemented by TIP4P/2005s yields larger (less negative) contrasts (Fig. 3C/D, yellow, blue, or red bars). Notably, owing to the small contrast ⎰ Δ*ρ*(**r**) d**r** of the overall XAO, Δ*R*_g_ values are highly sensitive with respect to water model-imposed variations of the hydration shell, leading to large Δ*R*_g_ variations of by up to 2 Å and even more (Fig. 4C/D, yellow, blue, and red bars; cf. Eq. 1).

### Hydration shell effect on *R*_g_ strongly depends on IDP conformation

To test how the peptide conformation influences the hydration shell of XAO, we computed hydration shell contrasts and Δ*R*_g_ values for 20 representative conformations of the aspartate mutant of XAO (Fig. S4) and analyzed two example conformations –one compact and one extended conformation– in detail in Fig. 6. We found that peptide conformations have only a small effect on the contrast (Fig. 6A/B) but may impose large variations of Δ*R*_g_ by up to ∼0.8 Å (Fig. 6C/D, Fig. S4). Notably, Δ*R*_g_ values neither correlate significantly with *R*_g_ values nor with *I*_0_, suggesting that subtle details of the peptide and hydration shell geometries determine the Δ*R*_g_ value.

**Figure 6.**
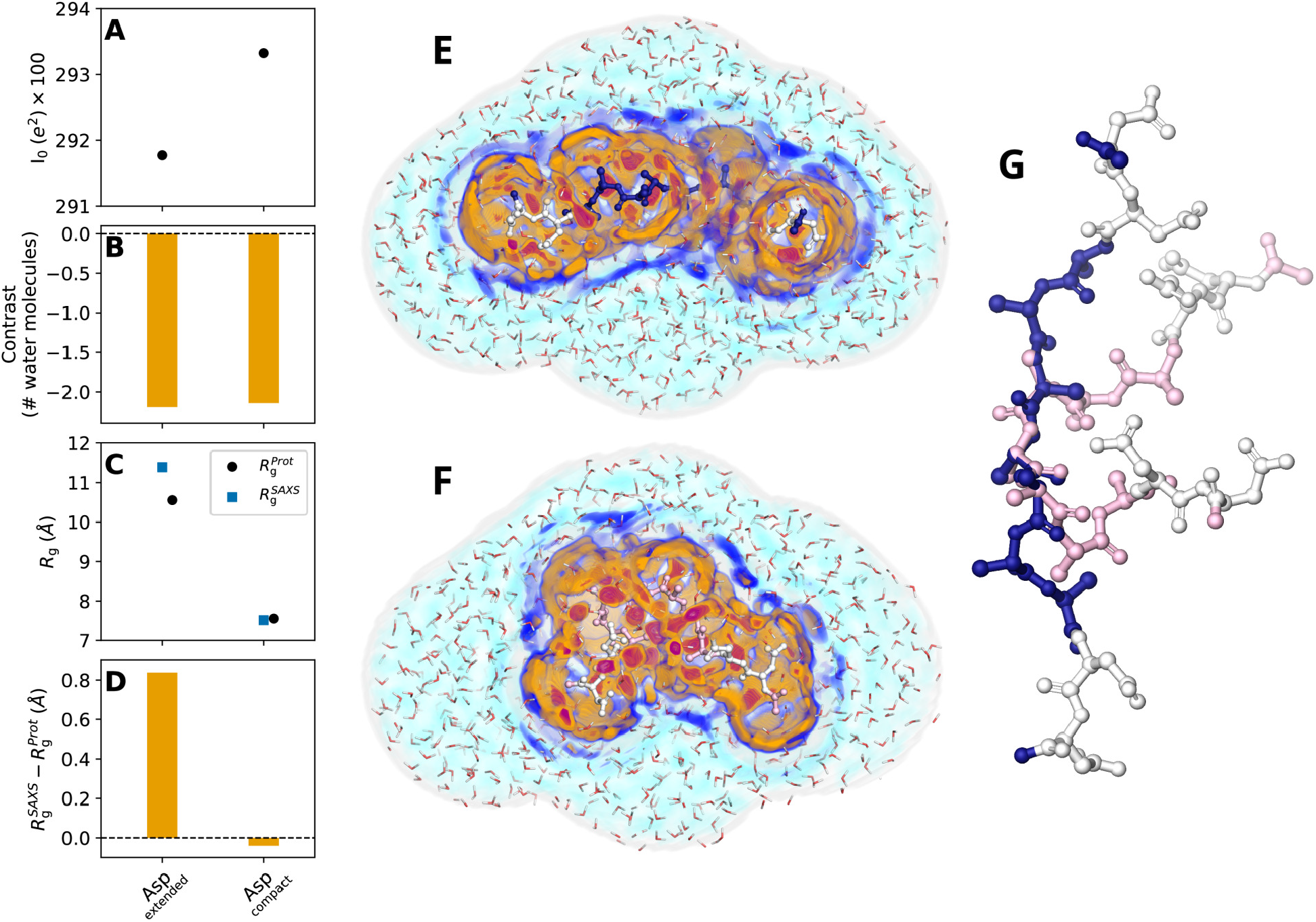
(A) Forward scattering *I*_0_, (B) hydration shell contrast, (C) *R*_g_, and (D) Δ*R*_g_ values of one extended and one compact conformation of the aspartate mutant of XAO obtained with TIP4P/2005 and ff03w. (E/F) Shaded colors show the three-dimensional solvent density maps from 50 ns MD simulations around the XAO mutant analyzed in panels A–D with color code taken from Fig. 1B/D. The density is overlayed with one MD frame, showing the XAO as balls/sticks and water molecules within the envelope as red/white lines. MD simulation used to compute density maps were carried out with restraints on all heavy atoms, thereby yielding spatially well-defined hydration shells. (G) Extended (blue/white) and compact conformation (pink/white) of XAO with four terminal residues mutated to aspartate (white).

### How surface-exposed chemical moieties determine the hydration shell contrast

The hydration shell contrasts computed for all proteinogenic amino acids (Fig. 3) enabled us to quantify how chemical modifications of surface-exposed moieties alter the hydration shell contrast. Table 1 lists eleven chemical modifications, together with the accompanying changes of contrasts 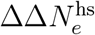 in number of electrons, as taken from the GB3 or XAO simulations with TIP4P/2005. Since one water molecule contains 10 electrons, the 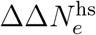 values may be translated to number of water molecules by dividing by 10. The values are qualitatively consistent among the analysis from GB3 and XAO. However, values differ quantitatively, likely reflecting that the degree to which these moieties are solvent-exposed differs between GB3 and XAO.

**Table 1.**
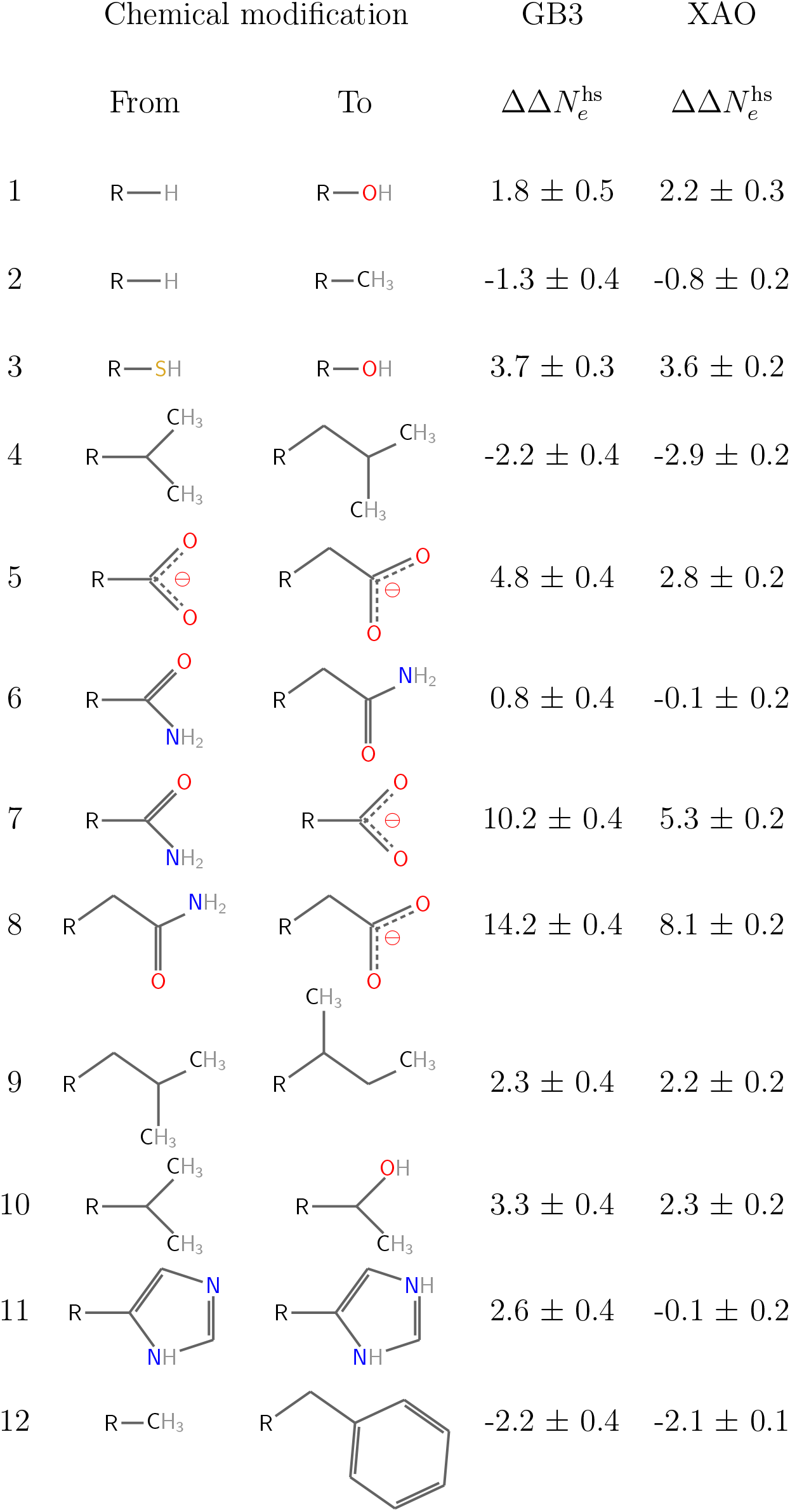
Change of hydration shell contrast in number of electrons upon various chemical modifications of surface-exposed amino acids. Errors denote 1 SE.

The analysis enables the following conclusions: (i) replacing a hydrogen (H) with a methyl group (CH_3_) decreases the contrast by ∼1 electron (Tab. 1, row 2). (ii) Replacing a methyl (CH_3_) with a hydroxyl group (OH) increases the contrast by ∼3 electrons (rows 1, 2, 10).

(iii) Replacing carbamoyl group (CONH_2_) as present in Asn or Gln with a carboxyl group (COO^−^) as present in Asp or Glu leads a marked increase of the contrast between 5 and 14 electrons (rows 7, 8). (iv) Extending a side chain by the addition of a CH_2_ group increases the contrast for a polar side chain (e.g. Asp→Glu, row 5) and decreases the contrast for apolar side chains (row 4), rationalized by the fact that longer side chains are more solvent-exposed. In other words, chemical modifications at the tip of longer side chains have a larger effect on the hydration shell contrast as compared to modifications at shorter side chains. Together, these values quantify how solvent-exposed chemical moieties modulate the electron density of the hydration shell of globular proteins or IDPs.

## Discussion

We quantified the effect of the 20 proteinogenic amino acids on the protein hydration shell density of the GB3 domain and the XAO peptide as representatives for the classes of globular or intrinsically disorders proteins with focus on two quantities encoded by SAXS curves: (i) the forward scattering *I*_0_, which reports on the overall contrast between the protein — including its hydration shell— relative to the buffer; and (ii) the radius of gyration *R*_g_, which is modified by the hydration shell relative to the *R*_g_ of the bare protein. Across both proteins and three different water models, we observed a consistent trend in hydration shell density at surface-exposed amino acids: anionic *>* cationic *>* polar *>* apolar residues. Substituting alanine with an anionic residue at the GB3 surface, the hydration shell density increased considerably and contained approximately 1 to 1.5 additional water molecules relative to the bulk density. Given that (i) water is nearly incompressible and (ii) amino acids are in contact with only a few water molecules, these values demonstrate a highly condensed packing of water at anionic amino acids and rationalize the presence of a marked hydration shell around anionic proteins.^20,49^ In contrast, replacing alanine with bulkier hydrophobic residues led to a decreased hydration shell density, which supports the presence of a water depletion layer previously reported for hydrophobic surfaces. ^36–38^

While the amino acid-specific effects on the hydration shell are qualitatively consistent between the GB3 domain and the XAO peptide, we also observed distinct differences in their hydration shells and their effects on SAXS curves. The hydration shell contrast was positive for the GB3 wild type and for most polar or charged GB3 mutants (Fig. 3A), indicating tightly packed water on the protein surface. In contrast, the hydration shell of most XAO variants revealed a negative contrast, indicating the presence of water depletion layers at the XAO surface (Fig. 3C). Thus, hydration shells of IDPs may differ substantially from hydration shells of globular proteins.

Furthermore, our calculations revealed that the effect of the hydration shell on *R*_g_ strongly depends on the conformation of XAO. This finding can be rationalized by the fact that the *R*_g_ of XAO is sensitive to the spatial distribution of hydration shell contrast: contrasts located farther from the XAO center of mass have greater effect on *R*_g_ as compared to contrasts closer the center of mass. Thus, the structure of the hydration shell and its effects on SAXS curves is controlled by an interplay between amino acid composition and peptide conformation.

Computing the hydration shell contrast from *I*_0_ values requires a definition of the protein volume or, equivalently, of the protein–water dividing surface. Indeed, different conventions for the dividing surface, for instance based on different Voronoi tessellation schemes, have led to slightly different protein volumes and different estimates for the hydration shell density.^45,59–62^ This unavoidable degree of ambiguity may explain why a previous study found similar water volumes at anionic, cationic, and polar moieties,^45^ whereas our analysis suggested by far larger contrasts (i.e., smaller water volumes) imposed by the anionic Asp/Glu residues. Critically, the pronounced hydration shells by Asp/Glu are confirmed by our Δ*R*_g_ calculation, which do not require assumptions on the dividing surface. In addition, they align with previous experimental SAXS/SANS studies of super-charged GFP variants and with large Δ*R*_g_ values found for the highly anionic glucose isomerase.^42,63^

We anticipate that the residue-resolved hydration shell contrast scores derived for all 20 proteinogenic amino acids will be useful for several future developments. Contrast scores may be used to parameterize computationally efficient SAXS curve predictions that account for residue-specific hydration while avoiding the need for explicit-solvent MD simulations for each protein conformation. Thereby, our calculations may bridge the gap between accurate yet computationally expensive explicit-solvent SAXS calculations^46,64–69^ and simplified implicit solvent methods that require fitting of the hydration shell against experimental data.^70,71^ In addition, quantifying residue-specific hydration will be key to understanding how targeted modulation of the water structure by protein–water interactions promotes biomolecular function, for instance in contexts of anti-freeze proteins, molecular recognition, or biomolecular phase separation. ^72–74^

## Methods

### Simulation setup and parameters for the GB3 domain

The initial structure of the GB3 domain was taken from the protein data bank (PDB;^75^ code: 1IGD^76^). Ten amino acids on the surface of the GB3 domain were selected and mutated to one of the 21 proteinogenic amino acid, involving two protonation states of histidine, with the software Chimera,^77^ namely residues 15, 18, 20, 22, 24, 27, 33, 37, 47, and 51 (Fig. 1A). Hydrogen atoms were added with pdb2gmx. MD simulations of GB3 were carried out with GROMACS, version 2020.3.^78^ Interactions of the proteins were described with the following variants of the Amber03 force field:^52^ ff03*,^79^ ff03w,^47^ and ff03ws.^51^ The starting structures were placed in a dodecahedral box, where the distance between the protein and the box edges was at least 2.0 nm, and solvated in TIP3P,^50^ TIP4P/2005,^48^ or TIP4P/2005s^51^ water. The simulation systems were neutralized by adding Na^+^ or Cl^−^ ions. After 400 steps of minimization with the steepest decent algorithm, the systems were equilibrated for 100 ps with harmonic position restraints applied to the heavy atoms of the proteins (force constant 1000 kJ mol^−1^nm^−2^). Subsequently, production runs were started for 50 ns with harmonic position restraints applied to the backbone atoms of the proteins (force constant 2000 kJ mol^−1^nm^−2^). The equations of motion were integrated using the leap-frog algorithm.^80^ The temperature was controlled at 298.15 K, using velocity rescaling (*τ* =1 ps).^81^ The pressure was controlled at 1 bar with the Berendsen thermostat (*τ* =1 ps)^82^ and with the Parrinello-Rahman thermostat (*τ* =5 ps)^83^ during equilibration and production simulations, respectively. The geometry of the water molecules was constrained with the SETTLE algorithm,^84^ and LINCS^85^ was used to constrain all other bond length. A time step of 2 fs was used. Dispersive interactions and short-range repulsion were described by a Lennard-Jones potential with a cut-off at 1 nm. The pressure and the energy were corrected for missing dispersion corrections beyond the cut-off. Neighbor lists were updated with the Verlet scheme. Coulomb interactions were computed with the smooth particle-mesh Ewald (PME) method.^86,87^ We used a Fourier spacing of approx. 0.12 nm, which was optimized by the Gromacs mdrun module at the beginning of each simulation.

### Simulation setup and parameters for the XAO peptide

To obtain an ensemble of XAO, we carried out maximum-entropy ensemble refinement^57^ of XAO against experimental SAXS data. To this end, four parallel replicas of XAO simulations were coupled to SAXS data as described in the Supplementary Information. From the ensemble, we obtained 20 frames that reasonably represent the confrontational space adopted by XAO, thus including compact and extended conformations. The simulation systems for the XAO conformations and mutants were setup as described above for the GB3 domain, except that the MD simulations were carried out with the GROMACS 2021.7.^78^ Side chains of ornithin (Orn) and 2,4-diaminobutyric acid (Dab) and were modeled based on the parameters for lysine by removing either one or two CH2 groups from the lysine topology, respectively. Four amino acids of the XAO, two Orn and two Dab two were mutated to one of the 21 proteinogenic amino acid with the software Chimera.^77^

### Explicit-solvent SAXS calculations

2251 simulation frames from the time interval between 5 and 50 ns from MD simulations of the GB3 domain and 2501 simulation frames from the time interval between 0 and 50 ns from simulations of the XAO peptide were used for SAXS calculations. The SAXS calculations were performed with GROMACS-SWAXS, an in-house modification of Gromacs 2021.7, as also implemented by the web server WAXSiS.^39,40,46^ The code and tutorials are available at https://cbjh.gitlab.io/gromacs-swaxs-docs/. For more background on explicit-solvent SAXS calculations we refer to recent reviews. ^41,88^ A spatial envelope was built around all solute frames from the proteins. Solvent atoms inside the envelope contributed to the calculated SAXS curves. The distance between the protein and the envelope surface was at least 9 Å, such that all water atoms of the hydration shell were included. The buffer subtraction was carried out using 2251 simulations frames of a pure-water simulation box, which was simulated for 50 ns and large enough to enclose the envelope. The orientational average was carried out using 200 **q**-vectors for the GB3 domain and 50 **q**-vectors for the XAO peptide for each absolute value of **q**, and the solvent electron density was corrected to the experimental value of 334 *e*/nm^3^ as described previously.^39^

Statistical errors were computed for simulations with the GB3 domain by binning the trajectory into 10 time blocks of 4.5 ns and computed the standard error. In the case of the XAO peptide, statistical errors were calculated from simulations of 20 independent conformations.

### Calculation of the hydration shell contrast

The forward scattering intensity 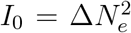 of a SAXS curve is given by the square of the contrast 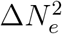 between solute (including the hydration shell) and the solvent in number of electrons. Thus, *I*_0_ follows by

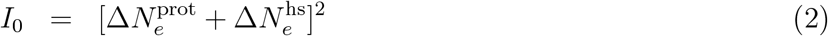

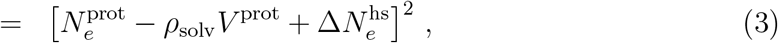

where 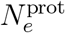 is the number of electrons of the protein, *V* ^prot^ the protein volume, 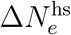 the contrast imposed by the hydration shell, and *ρ*_solv_ the solvent electron density taken as 334 *e*/nm^3^. Thus, we have

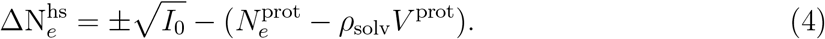

Here, the plus and minus signs correspond to the cases where 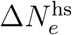 is positive or negative, respectively. In our implementation, 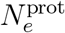 is taken from atomic form factor at zero scattering angle as defined via the Cromer-Mann parameters of the atoms.^89^ The volume of the solute *V* ^prot^ was defined as the cavity volume calculated with the 3V volume calculator^90^ with a grid spacing of 0.1 Å and a probe radius of 1.4 Å corresponding to the van der Waals radius of a water molecule. Volumes of GB3 and XAO variants are shown in Fig. S7.

## Acknowledgement

We thank Jan Lipfert for stimulating discussions and for sharing SAXS data of XAO. This study was supported by the Deutsche Forschungsgemeinschaft (DFG, German Research Foundation) via grants HU 1971/3-1 and INST 256/539-1.

## Supporting Information Available

Computational details for ensemble refinement of the XAO peptide and Figs. S1–S7.

