## Supplementary PDF for "Hydration shells of globular and intrinsically disordered proteins: effects of amino acid composition, peptide conformation, and force fields"

### Supporting Information Methods

#### Maximum-entropy refinement with the XAO peptide ensemble against SAXS data

To obtain a set of 20 XAO conformations that are representative for the XAO solution ensemble, we carried out SAXS-restrained ensemble simulations with commitment to the maximum entropy principle.[1] Four parallel XAO simulation replicas were coupled on-the-fly to the SAXS curve taken from Ref. 2. Simulations we carried out with GROMACS-SWAXS, version 2021.5, as freely available at https://gitlab.com/cbjh/gromacs-swaxs. Documentation for GROMACS-SWAXS is available at https://cbjh.gitlab.io/gromacs-swaxs-docs.

Four starting structures for the SAXS-restrained simulations were taken from the XAO ensemble refined against NMR data by Makowska et al.[3] and set up for simulations as described in the Methods. To couple the simulations to the SAXS data, SAXS curves were computed from the simulations on-the-fly using explicit-solvent SAXS calculations, thereby taking scattering contributions from the hydration shell into account.[4–6] SAXS curves were averaged on-the-fly using a memory kernel that decays exponentially into to the past using a memory time of 100 ps [molecular dynamics parameter (mdp) option `waxs-tau`]. A $q$ range from 0.065 Å$^{-1}$ to 0.58 Å$^{-1}$ with 30 equally-spaced $q$-point was used (mdp options `waxs-startq, waxs-endq, waxs-nq`). The SAXS curve was updated every 125 ps (mdp options `waxs-nstcalc` together with `dt`). A force constant of unity was applied, and the restraints were turned on gradually over 10 ns (mdp options `waxs-fc, waxs-t-target`). During the simulations, and prior to computing SAXS-derived forces, the experimental SAXS curve was fitted to the calculated curve via $I_{exp,fit}(q) = f \cdot I_{exp} + c$, by minimizing $\chi^2$ with respect to the calculated curve. Here the factor $f$ accounts for the overall scale, and the offset $c$ accounts for a putative uncertainty from the buffer subtraction. No fitting parameters owing to the hydration layer or excluded solvent were used, implying that also the radius of gyration was not adjusted by the fitting parameters. The agreement of the SAXS curve obtained from the refined XAO ensemble with the experimental data is shown in Fig. S1.

SAXS-restrained simulations were carried out for 150 ns. The spatial envelope was built at a distance of 12 Å from all XAO atoms during free simulation that started from four different structures. Solvent atoms within the envelope contributed to the calculated SAXS curve as described previously.[5] The temperature was controlled at 298.15 K using a stochastic dynamics integrator.[7] All other simulation parameters were chosen as described in the Methods.

From the four trajectories collected from the four parallel replicas, five configurations each were taken from the simulation times 30 ns, 60 ns, 90 ns, 120 ns and 150 ns, thereby providing 20 independent conformations. These conformations were mutated as described in the main text and used for follow-up SAXS calculations. During follow-up simulations, these

20 conformations were maintained by applying positions restraints either to the backbone or to heavy atoms, as described above.

### Supporting Information Figures

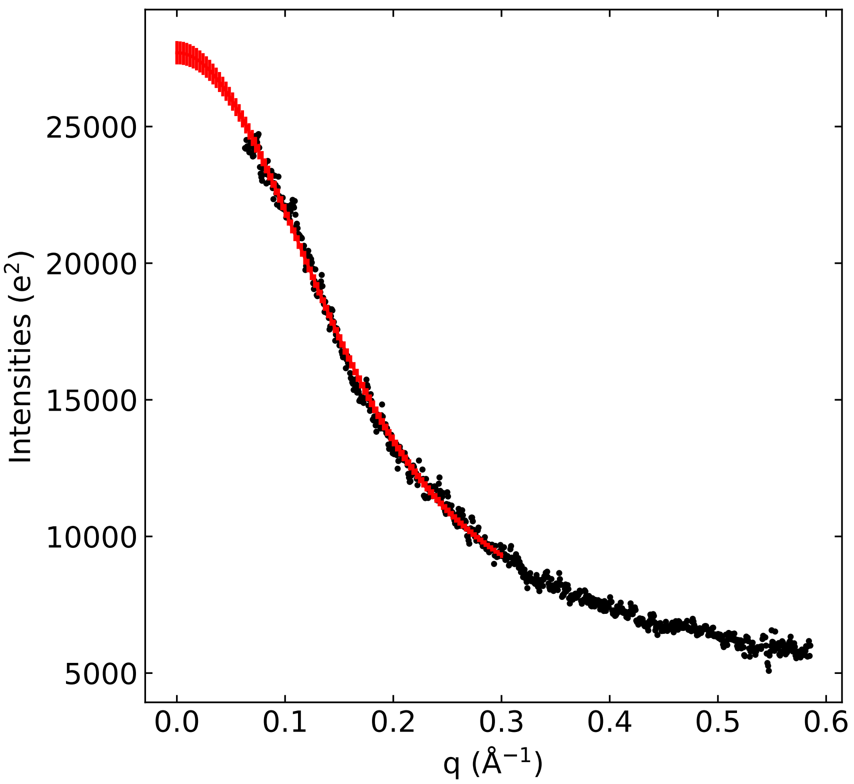


Figure S1: Experimental SAXS data by Zagrovic *et al.*[2] (black dots) and SAXS curve of XAO ensemble obtained by maximum-entropy ensemble refinement.[1] From the refined XAO ensemble, 20 frames were selected as representative conformations of the heterogeneous XAO ensemble and used subsequently for computing SAXS curves of XAO mutant.

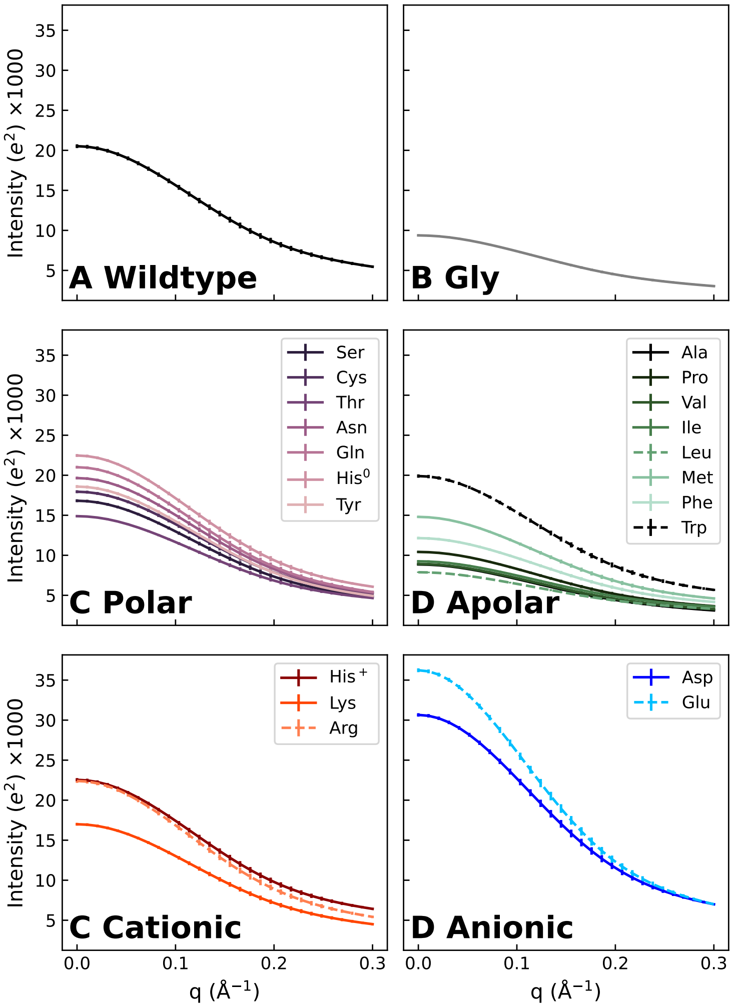


Figure S2: SAXS curves of the heterogeneous ensemble of the XAO peptide from explicit-solvent SAXS calculations with the TIP4P/2005 water model in combination with the ff03w protein force field. Backbone positions were restrained in simulations for all XAO mutants to the backbone positions of the XAO wild type ensemble refined against experimental SAXS data (see Fig. S1), suggesting that variations among the computed SAXS curves are purely caused by presence of four different amino acids (at fixed backbone positions) and by variations of the hydration shell. SAXS curves are shown (A) for the XAO wild type and (B–F) for for 21 mutants with four mutated surface-exposed amino acids each (for color code and line style, see legends). For clarity, SAXS curves are grouped by the amino acid property (glycine, polar, apolar, cationic, anionic) in panels B–F.

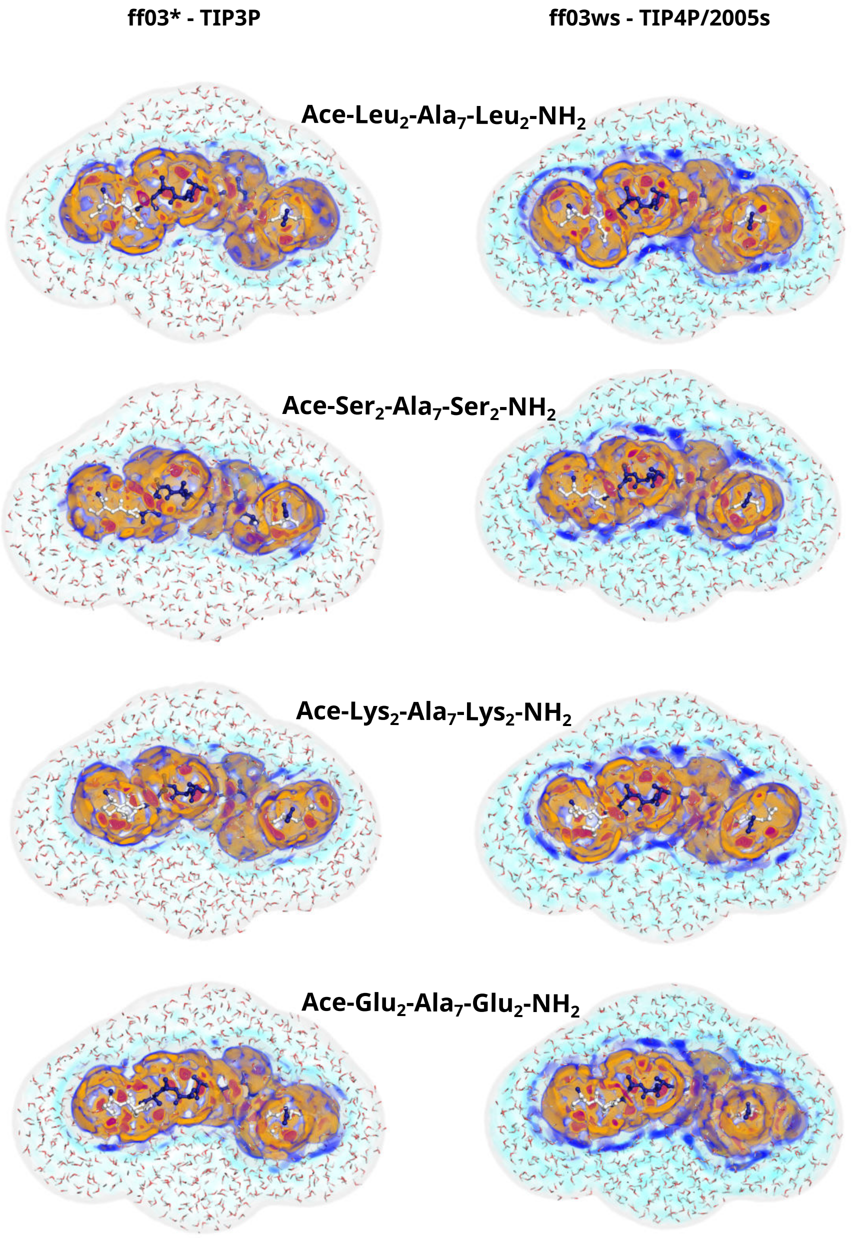


Figure S3: Three-dimensional densities of the hydration shell around the XAO mutants with four leucine, serine, lysine, or glutamate residues at the termini (see labels). The densities were calculated from simulations using ff03$^*$ in conjunction with TIP3P (left column) or using ff03ws in conjunction with TIP4P/2005s (right column). Color code is taken from Fig. 1B/D. The solvent densities depend on amino acid type and on the force field.

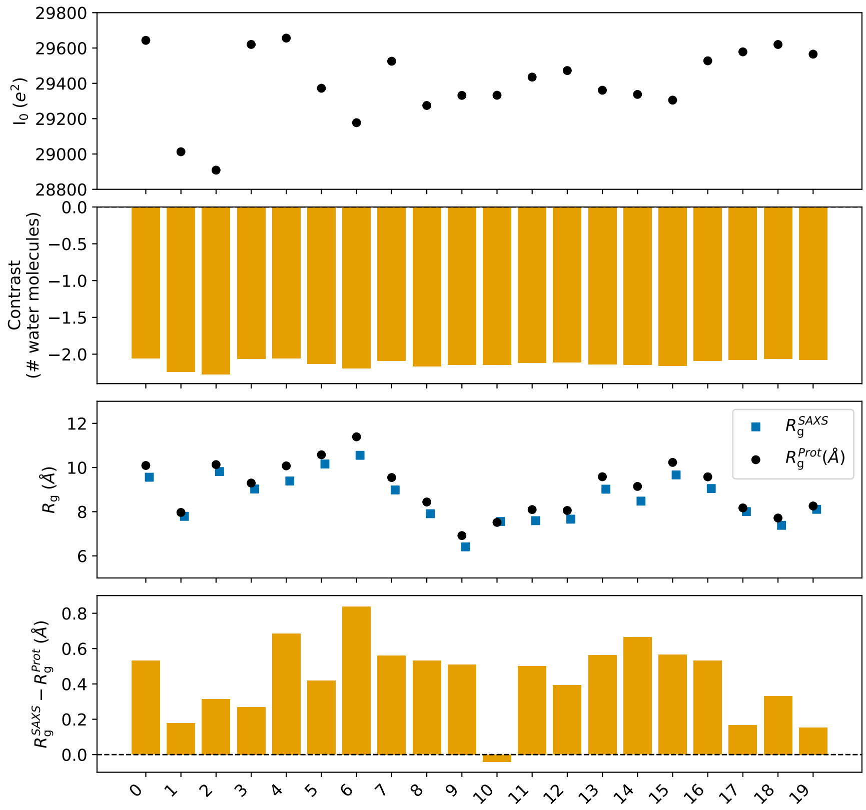


Figure S4: Forward scattering $I_0$, hydration shell contrast in number of water molecules, $R_g$, and $\Delta R_g$ values for 20 conformations of the aspartate mutant of XAO, obtained with TIP4P/2005 and ff03w. The different conformations impose similar hydration shell contrasts, yet lead to greatly different $\Delta R_g$ values.

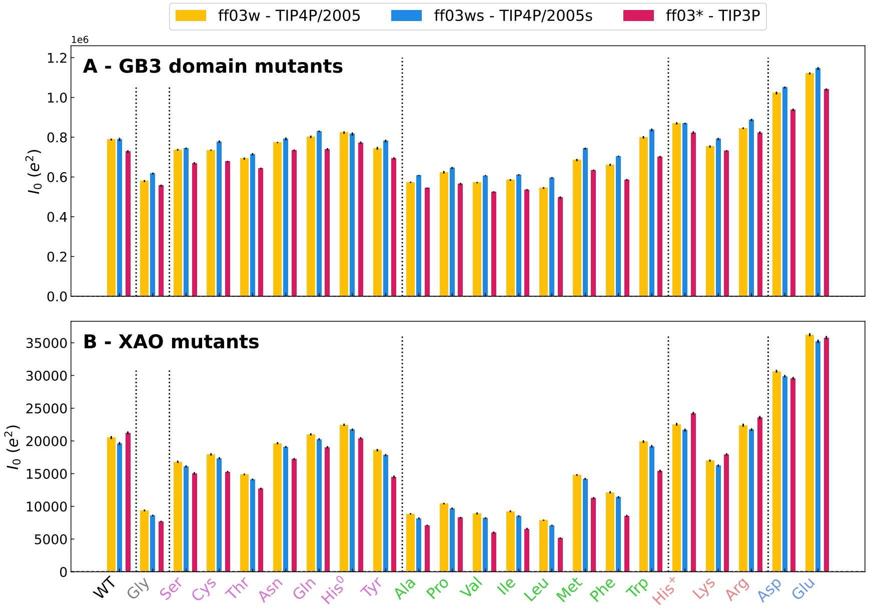


Figure S5: Forward scattering intensity $I_0$ from SAXS curves for wild type and 21 mutants (see labels on abscissa) of (A) GB3 domain and (B) XAO peptide from simulations with three different combinations of protein force field and water model (for color code, see legend).

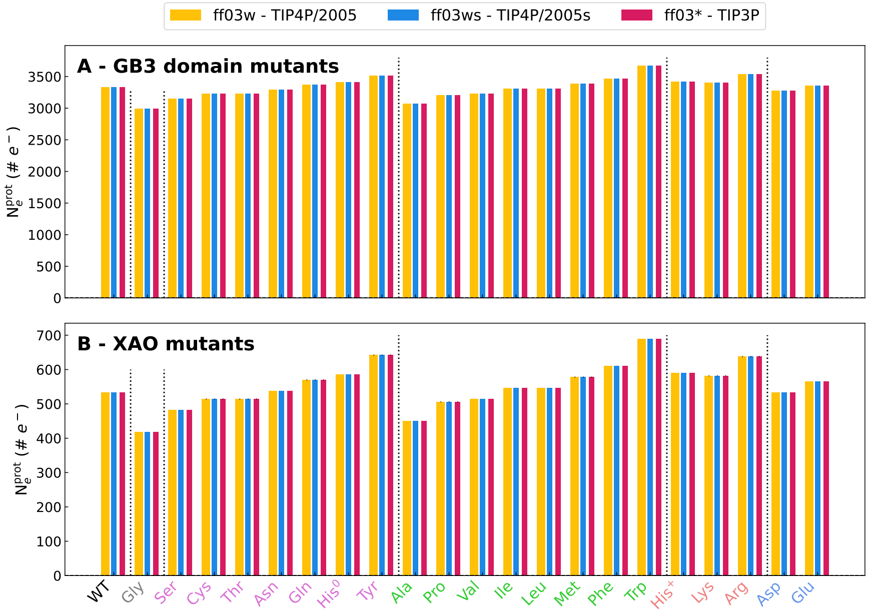


Figure S6: Number electrons (# $e^-$) of the solute for WT and 21 mutants of (A) GB3 domaind and (B) XAO peptide from simulations with three different combinations of protein force field and water model (for color code, see legend).

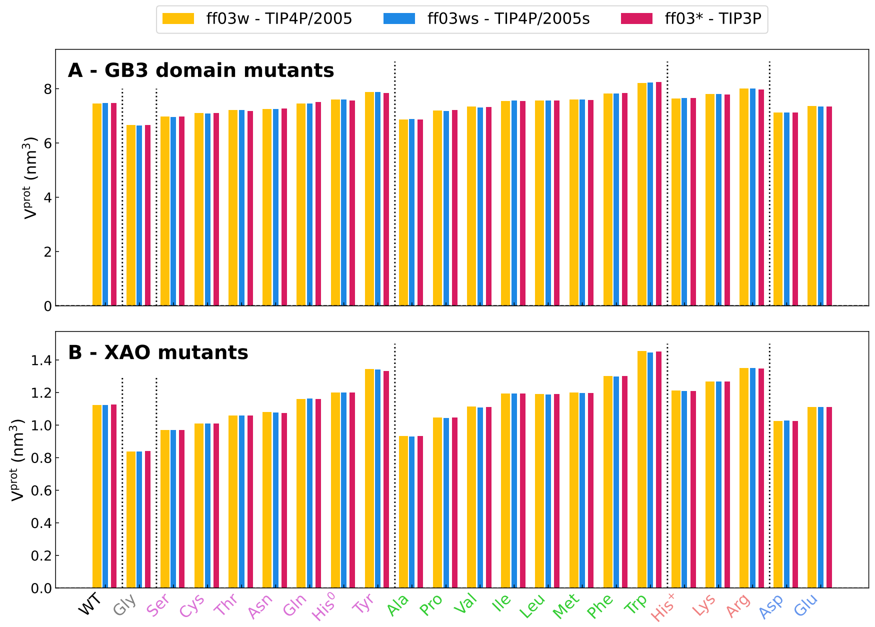


Figure S7: Volumes of WT and 21 mutants (see labels on abscissa) of (A) GB3 domain and (B) XAO peptide from simulations with three different combinations of protein force field and water model (for color code, see legend).
